# Previously Hidden Dynamics at the TCR-pMHC Interface Revealed

**DOI:** 10.1101/223628

**Authors:** James Fodor, Blake T. Riley, Natalie A. Borg, Ashley M. Buckle

## Abstract

A structural characterization of the interaction between T cell receptors (TCR) and cognate peptide-MHC (pMHC) is central to understanding adaptive T cell mediated immunity. X-ray crystallography, although the source of much structural data, traditionally provides only a static snapshot of the protein. Given the emerging evidence for the important role of conformational dynamics in protein function, we interrogated 309 crystallographic structures of pMHC complexes using ensemble refinement, a technique that can extract dynamic information from the X-ray data. We found that in a large number of systems ensemble methods were able to uncover previously hidden evidence of significant conformational plasticity, thereby revealing additional information that can build upon and significantly enhance functional interpretations that are based on a single static structure. Notable examples include the interpretation of differences in the disease association of HLA subtypes, the relationship between peptide prominence and TCR recognition, the role of conformational flexibility in vaccine design, and discriminating between induced fit and conformational selection models of TCR binding. We show that the currently widespread practise of analyzing pMHC interactions via the study of a single crystallographic structure does not make use of pertinent and easily accessible information from X-ray data concerning alternative protein conformations. This new analysis therefore not only highlights the capacity for ensemble methods to significantly enrich the interpretation of decades of structural data, but also provides previously missing information concerning the dynamics of existing characterized TCR-pMHC interactions.

## Introduction

Cytotoxic T cells express on their cell surface αβ T cell receptors (TCR), which engage endogenous peptide antigens from infected or malignant cells that are displayed by major histocompatibility complex (MHC) proteins. Productive engagement of the TCR with the peptide-MHC complex (pMHC) leads to T cell activation and intracellular signal transduction via the CD3 signaling subunits. This TCR-pMHC molecular recognition event is key to the cell-mediated adaptive immune response, and has therefore been widely studied using X-ray crystallography. These studies have focused on the conformation of the peptide when bound to the MHC and the specific interactions between peptide and MHC side-chains, with or without bound TCR. While providing insights into the key features of TCR-pMHC recognition, the detailed examination of a single static structure by X-ray crystallography reveals little about the conformational dynamics and degree of flexibility of the system in solution^1-3^. Given that protein flexibility is inextricably linked to protein function, consideration of protein dynamics is essential in order to achieve a full understanding of pMHC and TCR-pMHC binding. Recent years have seen an increasing use of atomistic molecular dynamics (MD) simulations, which can probe the conformational flexibility of pMHC systems by computing iterative numerical solutions of Newton’s equations of motion as the system evolves over time ^4^. Unfortunately, the heavy computational demands of MD limits the technique to examination of relatively short time spans (usually less than a microsecond), and thereby excludes many larger motions that occur over longer, more biologically relevant timescales. Results of MD are also somewhat difficult to validate given the known imperfections of existing force fields ^5-7^.

Ensemble X-ray structure refinement is a refinement technique that can be employed to extend the interpretation of crystallographic data, and obtain information about protein dynamics without the necessity of running expensive MD simulations ^8^. Instead of the usual practice of refining against a set of crystallographic reflections to produce a single protein structure, ensemble refinement combines the experimentally-derived reflection data with the results of a short, steered MD simulation to produce an ensemble of similar but distinct conformations of the protein. The structural diversity observed within the resulting ensemble thus provides information about the conformational flexibility of the protein, and in many cases improves the agreement between the structure and data as measured by the crystallographic free R-factor (*R*_free_). Notable recent applications of this method include an investigation of the conformational landscape of an insect carboxylesterase protein over the course of its catalytic cycle ^9^, and the discovery of a self-inhibition mechanism mediating the transition between active and inactive protein of human factor D ^10^. Ensemble refinement is therefore a technique for extracting dynamical information already present in X-ray reflection data but overlooked in the process of producing a single static structural model. As such, we hypothesized that when applied to pMHC and TCR-pMHC systems, ensemble refinement would uncover evidence of molecular motion that has hitherto been hidden. We further hypothesized that the dynamic information revealed would in many cases significantly enhance existing single structure analyses of these systems. Here we test these hypotheses by applying ensemble refinement to investigate protein dynamics at the TCR-pMHC interface.

## Results

To acquire a sample of appropriate pMHC systems whose dynamics could be investigated using ensemble refinement, we first searched the Protein Data Bank (PDB) for pMHC complexes with a resolution of better than 3 Ångströms. Some of the resulting entries were rejected owing to lack of the necessary crystallographic reflection files, leaving 284 pMHC structures which were subjected to ensemble refinement as described in Methods. Each of these systems consisted of the heavy and light chains of the MHC molecule, the bound peptide, and also the α and β chains of the TCR when this was present. The ensemble refinement process takes as input a single crystallographic structure and returns as output an ensemble of up to several dozen structures that collectively fit the X-ray diffraction data roughly as well as the original single structure. Of all the systems we subjected to this process, we informally estimated that about half contained conformations noticeably different from the original PDB structures. It was therefore necessary to reduce this to a more manageable number of systems that we could subject to detailed structural analysis. First, in order to eliminate possible confounds we discarded all systems with bound ligands and missing peptide residues. Next, we ranked these systems according to the degree of conformational flexibility displayed by the peptide and MHC peptide-binding helices, as measured by the root mean square fluctuation (RMSF) of these residues calculated over the ensemble structures. To further focus our analysis, we then considered only published structures of human class I MHCs (human leukocyte antigens or HLA), selecting the top 30 most flexible ensembles from the RMSF ranking as being the most relevant for testing our hypothesis. For each of these systems, we compared our results in detail to those reported in each original publication.

In comparing our ensemble results with the analyses reported in the original papers, we found it necessary to run an additional 25 ensembles to properly interpret our ensemble results, thus raising the total number of systems for which ensembles were calculated to 309. Given space limitations, from these 30 systems we selected 12 illustrative cases in which our ensemble results provided the greatest functional insights pertinent to the claims made in the original literature. Since in some cases the original study compared multiple pMHC systems, these 12 examples included a total of 20 distinct PDB structures, all of which are summarized in Table 1. In our analysis of these systems, we highlight several new insights into TCR-pMHC engagement yielded by ensemble analysis compared to traditional approaches that consider only a single static structure. To supplement our ensemble results, five of the systems were selected for analysis in standard all-atom MD simulations, and the results compared to the corresponding ensembles.

**Table 1.**
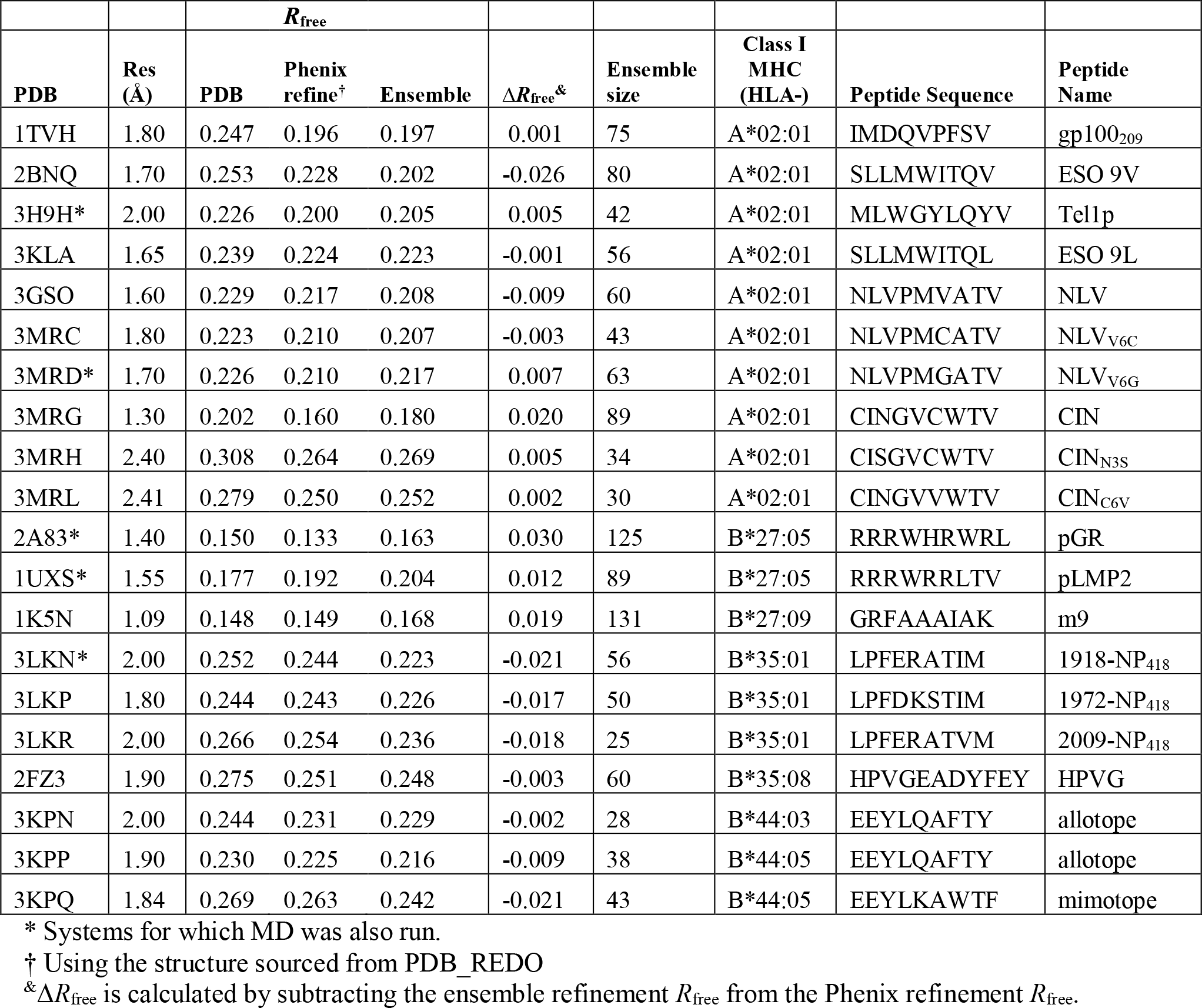
pMHC systems investigated by ensemble refinement.

### Conformational polymorphism and peptide variants

A common goal in studying HLA-A*02 systems is to determine the structural basis for differential TCR recognition of peptide or MHC variants, and the effect of such variations on T cell activation. For example, Chen et al^11^ compared two peptide analogs of the NY-ESO-1 tumor antigen peptide, ESO 9L and ESO 9V, both bound to HLA-A*02:01. The peptides are generally very similar, except that HLA-A*02-ESO 9V has a six-fold greater binding affinity for the 1G4 TCR compared to HLA-A*02-ESO 9L. The authors attribute this affinity difference to a shift in the position of residues P4-P9 of the ESO 9L peptide relative to their location in the ESO 9V peptide (Figure 1A). This change in position is in turn attributed to the greater length of the P9-Leu residue compared to P9-Val, which prevents the ESO 9L peptide from sitting as low in the binding grove as ESO 9V. Furthermore, the I6 residue (found in both ESO 9L and ESO 9V) was identified as blocking a possible conformational switch involving alpha helix residues Tyr116 and Arg97^12^ (Figure 1A, coral/steel blue). Specifically, the rotation of Tyr116 would free sufficient space to allow the longer P9-Leu residue to fit in the groove, however this rotation is blocked by the orientations of Arg97 and I6. Our ensemble results enrich this static picture by showing that the ESO 9L peptide is in fact easily flexible enough for the ‘forbidden’ Tyr116-Arg97 flip to occur, thereby enabling the peptide to adopt a wide range of conformations not seen in the original static structure (Figure 1B). This indicates that a full explanation of the difference in affinity of the 1G4 TCR for HLA-A*02-ESO 9V versus HLA-A*02-ESO 9L will incorporate their dynamic flexibility as a constraint on the interpretation of observed static structural differences. Similar bond flips have been observed in other ensembles of HLA-A*02:01 systems. Borbulevych et al^12^, for instance, found that a melanoma antigen (gp100_209_) bound to HLA-A*02:01 in two distinct conformations, differing in a bond flip of one key side chain (P7-Phe) (Figure 1C). Our ensemble results support the existence of both of these alternate conformations, though the large number of additional conformations observed highlight that for these systems it may be more accurate to speak of conformational polymorphism than conformational dimorphism (Figure 1D).

**Figure 1:**
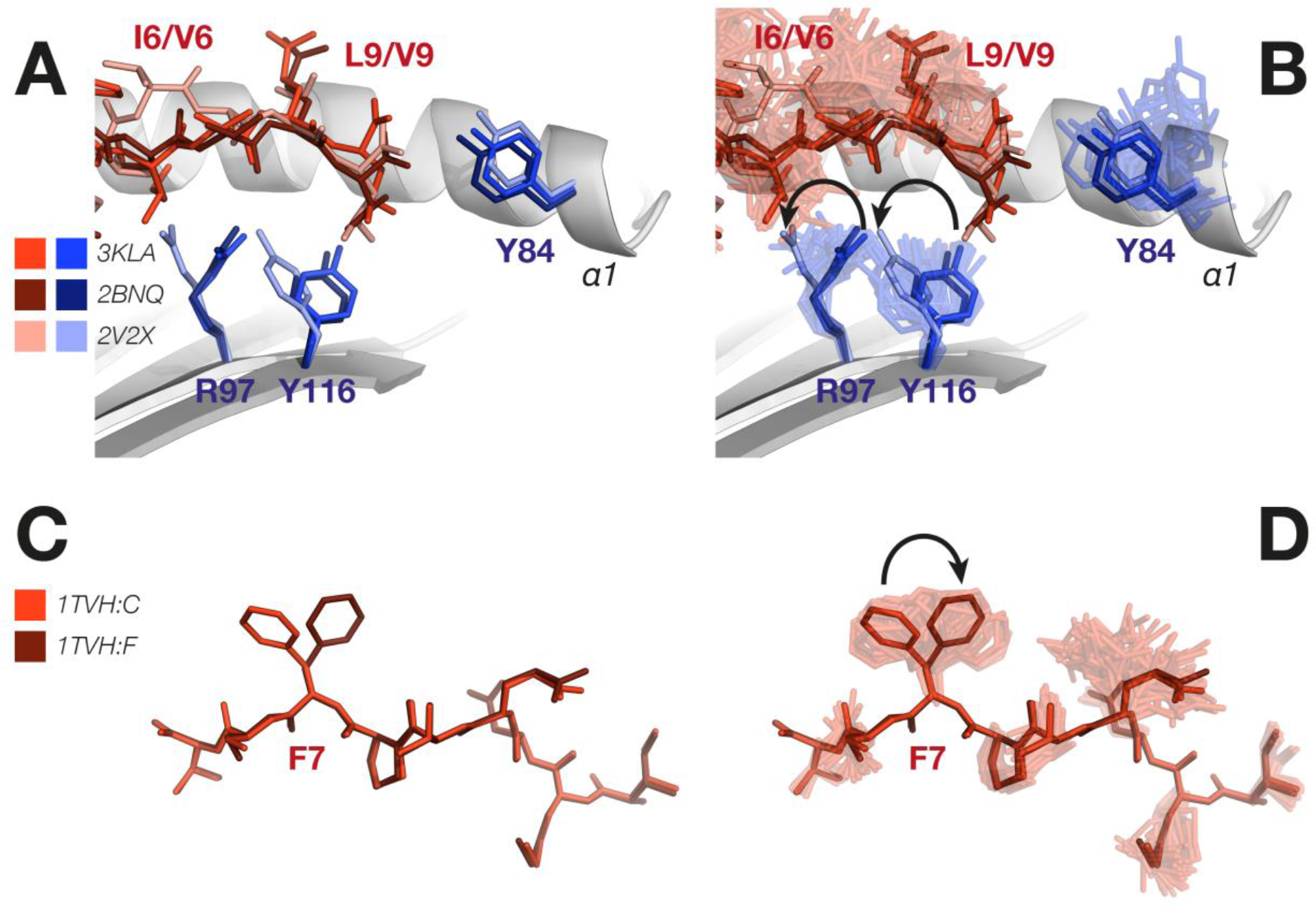
Conformational polymorphism and peptide variants. **A.** An overlay of HLA-A*02:01-ESO 9L (red/blue) (PDB ID 3KLA) and HLA-A*02:01-ESO 9V (brown/navy) (PDB ID 2BNQ), and an alternate ‘forbidden’ conformation of a similar peptide (SLFNTVATL, pale red/pale blue) bound to HLA-A*02:01 (PDB ID 2V2X), compared to **B.** ensemble results of HLA-A*02:01-ESO 9L showing high flexibility of the peptide, and MHC side chains undergoing the ‘forbidden’ bond flips. The MHC α2-helix is omitted. **C.** Two conformations of a melanoma antigen (gp100_209_) observed in different molecules of the crystallographic asymmetric unit (PDB ID 1TVH, chains C and F), and **D.** ensemble results from a single molecule showing interconversion between these conformations, in particular the side chain rotation observed in P7-Phe. MHC α-helices have been omitted.

### Conformational populations of pMHC

As in the case of HLA-A*02 systems, analysis of HLA-B*27 pMHC systems is also enriched by the use of ensemble methods to highlight temporal variation in peptide conformation. Fiorillo et al^13^, for example, compared the structure of HLA-B*27:05 and HLA-B*27:09 bound to the Epstein-Barr virus (EBV)-latent membrane protein 2 (termed pLMP2). These two alleles differ only by a single residue, with Asp116 in HLA-B*27:05 substituted with His116 in HLA-B*27:09. The authors observed a substantial difference in the conformation of the peptide between the two alleles (Figure 2A), which they argue may contribute to the observed differential TCR recognition. Ensemble results for these systems show that, while the two peptide conformations are not observed to fully interconvert, the peptide side chains are highly flexible, especially around the P4-P7 residues where the largest conformational difference is observed (Figure 2B). Our ensemble analysis thus renders visible an aspect of the two peptides that is hidden in a static analysis, namely that their conformational populations may show considerable overlap even when individual structures differ considerably. This is particularly relevant when considering that peptide flexibility at physiological temperatures will almost certainly exceed that observed at cryogenic temperatures used in crystallographic data collection. Further support for this is observed in our MD results for the HLA-B*27:09 system, which show a similar pattern of flexibility to that observed in the ensembles, with even clearer evidence of interconversion between the two conformations (Figure S1A and S1B). Taken together, our ensemble and MD results indicate that at physiological temperatures, the two distinct conformations observed by Fiorillo et al. are most likely part of a large ensemble of conformations that the peptide adopts when bound to the MHC. Ensemble methods thus enhance the static picture of peptide structure by revealing how conformational populations in the pMHC are perturbed by polymorphisms, and how this altered energy landscape affects TCR engagement.

**Figure 2:**
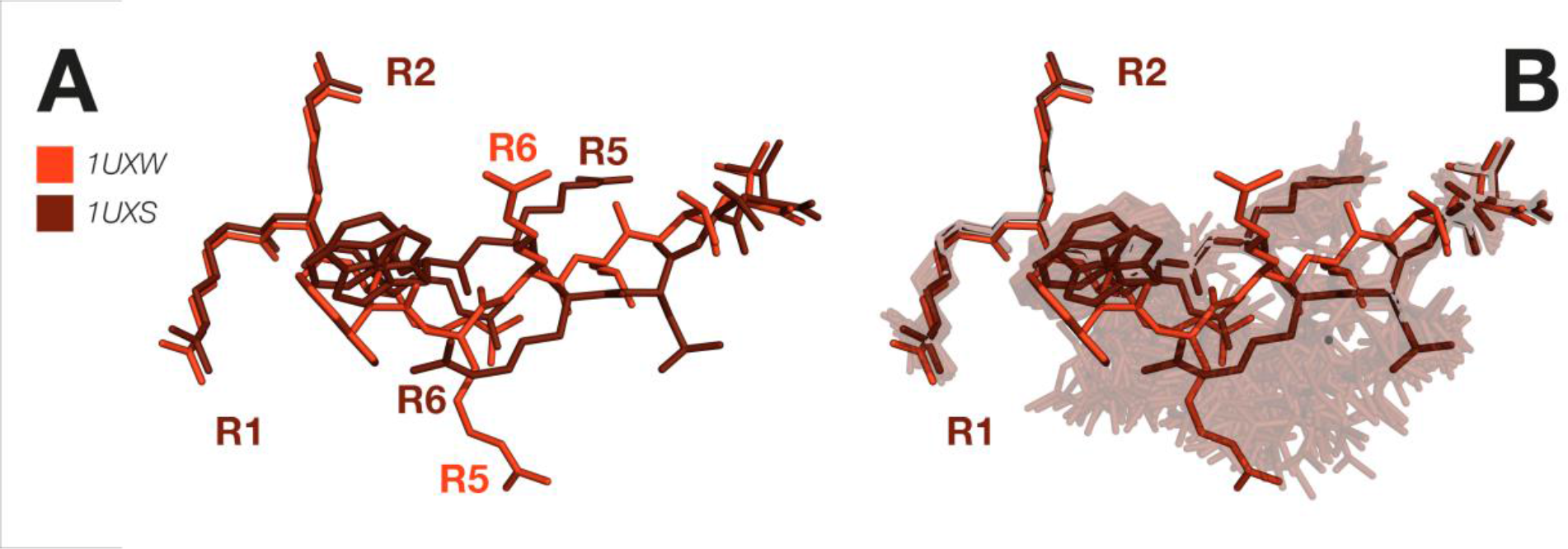
Conformational populations in MHC alleles. **A.** Superposition of original structures of Epstein-Barr virus pLMP2 peptide bound to HLA-B*27:05 (red) (PDB ID 1UXW) and HLA-B*27:09 (brown) (PDB ID 1UXS) showing the different conformations. **B.** Ensemble results of the HLA-B*27:09 system, showing the degree of flexibility and possibility of interconversion of the two binding modes. MHC α-helices have been omitted.

### Interpretation of static interactions

Ensemble methods can build upon traditional methods for inferring the functional relevance of networks of bonding interactions that are observed in static pMHC structures. For example, it has been proposed that the reason why HLA-B*27:05, but not HLA-B*27:09, is associated with spondyloarthropathies^15^ is because these two alleles elicit different T cell responses to arthritogenic peptides. To determine whether structural differences could account for this allele-specific disease association, Hülsmeyer et al^14^ compared the structures of HLA-B*27:05 and HLA-B*27:09, each bound to the same token peptide (m9). In their analysis they identify several subtle structural differences between the F-pocket regions of the two alleles, related to the exact position and orientation of P9-Lys, several of the surrounding MHC side-chains, and the network of water molecules mediating salt bridges between these residues (Figure 3A). They suggest that such differences alter the dynamics of the HLA-B*27:09 F-pocket region and accordingly could affect TCR recognition. Our ensemble analysis indicates that the F-pocket region of HLA-B*27:09 is highly flexible, with all of the indicated side-chains displaying considerable variation in position and orientation (Figure 3B). Given this substantial intrinsic flexibility, it is likely any hydrogen bonds revealed from a single static structure exist only transiently. Ensemble methods thereby expand on what can be concluded from a single static structure, which by design shows only a single conformation of the system as it exists at a single moment in time. Further work is necessary to determine how these differing patterns of side-chain and peptide conformational variability can account for the differential disease association of these alleles.

**Figure 3:**
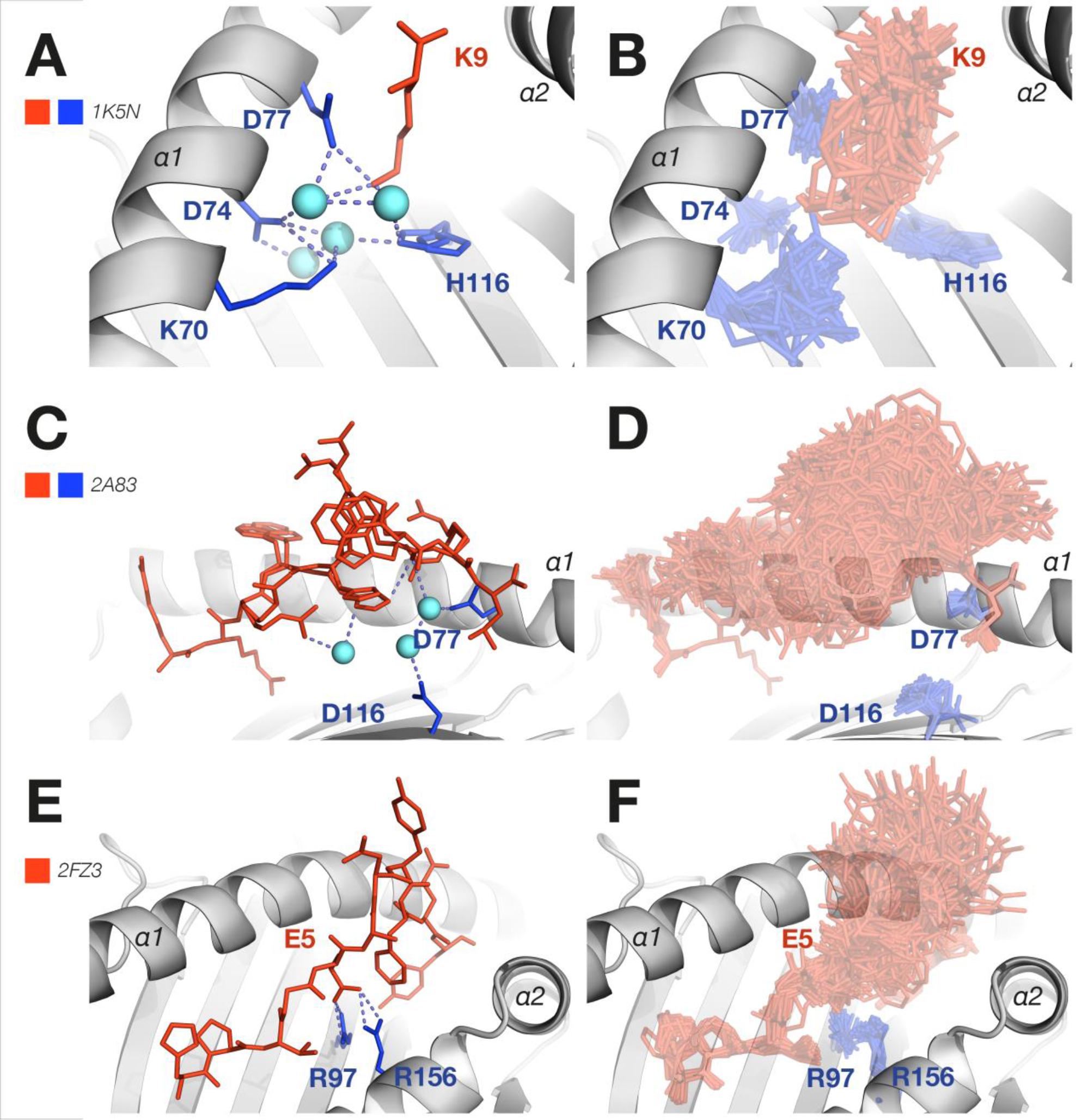
Interpretation of static contacts. **A.** Original structure of HLA-B*27:09-m9 (PDB ID 1K5N) with proposed stabilizing bonding network of the F-pocket region, compared to **B.** ensemble results, illustrating how many of the hydrogen bonds (shown in dashed lines) identified in the original structure are not stable over the ensemble. Water molecules are shown as light blue spheres. MHC α-helices have been omitted. **C.** In the original structure two conformations of the glucagon-derived self-peptide were observed when bound to HLA-B*27:05 (PDB ID 2A83) showing the proposed conserved hydrogen-bonding network. Hydrogen bonds are depicted as dashed lines, and water molecules are shown as light blue spheres. **D.** Ensemble results for the same system showing the hydrogen-bonding network is not conserved over the ensemble. The MHC α2-helices have been omitted. **E.** Original structure (PDB ID 2FZ3) of Epstein-Barr virus peptide HPVG bound to HLA-B*35:08 which the authors describe as ‘relatively rigid’, compared to **F.** the ensemble results of the same system showing considerable conformational variability of this peptide.

In a related study, X-ray crystallography was used to examine the association of HLA-B*27:05 with spondyloarthropathies^15^, and investigate the potential role of molecular mimicry and T cell cross-reactivity in pathogenesis more generally. The structure of HLA-B*27:05 bound to a glucagon-derived self-peptide pGR was determined. This pGR peptide was selected because of its sequence similarity to both the self-peptide pVIPR and the foreign peptide pLMP2, each of which cross-reacts with T cells in ankylosing spondylitis patients more often than in healthy controls. The aim of the study was to determine if pGR could adopt the structurally similar and unusual conformation observed of both pVIPR and pLMP2 when bound to HLA-B*27:05, thereby indicating that the observed cytotoxic T cell cross-reactivity may be due to molecular mimicry. The crystallographic structure contained two distinct conformations of pGR, along with a network of contacts supporting those conformations (Figure 3C). Both pGR conformations were also observed in our ensemble, as well as a wide range of intermediate conformations and further variants (Figure 3D). We also conducted MD on this system and found a very similar overall pattern of peptide flexibility to that seen in the ensembles (Figure S1C and S1D). Both our ensemble and MD results therefore indicate that many hydrogen-bonding networks in pMHC systems are not stable at physiological temperatures, but are in a constant state of dynamic flux.

To investigate the influence of MHC polymorphism on peptide conformation and the T cell response, the structures of HLA-B*35:01 and HLA-B*35:08 bound to the same EBV peptide (HPVG) were determined^16^ (Figure 3E). These two alleles differ only at position 156, with Leu156 found in HLA-B*35:01 compared with Arg156 in HLA-B*35:08. The authors identified a set of hydrogen bonds between the P5-Glu residue and two MHC side chains of HLA-B*35:08, which are not found in HLA-B*35:01. The authors attribute these hydrogen bonds as likely responsible for the ‘relatively rigid’ nature of the HPVG peptide when bound to HLA-B*35:08 (as determined by B factor analysis), compared to its very high flexibility when bound to HLA-B*35:01, and show that peptide mobility influences the T cell response. Our ensemble results reveal that HPVG too has considerable flexibility when bound to HLA-B*35:08 (Figure 3F), with the stabilizing interactions identified by the authors present only some of the time. This indicates that the stabilization produced by static bonding networks comes in varying degrees, and may still result in an overall highly flexible system.

### Prominence of MHC-bound peptides and TCR recognition

The degree to which MHC-bound peptides protrude from the binding cleft, and the effect this has on TCR recognition, is another type of structural analysis that can be augmented by the use of ensemble methods. For example, Reiser et al^17^ sought to identify pMHC structural features that correlate with pMHC immunogenicity by examining eight different viral and tumor peptides bound to HLA-A*02:01. While no linear relationship was found between peptide solvent exposure and the frequency of pMHC-specific T cells, several structural differences between the peptides were identified as likely causes of varying pMHC antigenicity, suggesting the existence of an optimal intermediate degree of prominence^17^. In particular, differences in the conformations of the wild type NLV peptide and the NLV_V6C_ and NLV_V6G_ peptide variants were attributed to the absence of the buried stabilizing P6-Val side chain in the modified peptides (Figure 4A). Our ensemble analysis shows that each of the modified NLV peptides (especially the V6G variant) exhibits considerable flexibility around the P5 and P6 residues (Figure 4A-D), which suggests that when intrinsic structural flexibility is considered these variants are more similar to each other than the static structures alone suggest. This finding is further supported by our MD simulation results of the NLV_V6G_ peptide system, which also showed considerable variation around these residues (Figure S1E and S1F). In a similar analysis of static structures of HLA-A*02:01 bound to the CIN, CINN3S, or CINC6V peptide variants (Figure 4E), it was found that the frequency of pMHC-specific T cells is influenced by peptide prominence from the MHC binding cleft, peptide mobility, and by the altered conformation of the MHC Gln155 side chain between the systems. Again, our ensembles show that both the peptide prominence and the position of the Gln155 residue vary considerably within each system (Figure 4F-H). This further emphasizes the value of ensemble methods in providing otherwise hidden information about peptide mobility, and illustrating that peptide prominence should be considered over a range of conformations in order to accurately characterize its potential relationship with TCR recognition.

**Figure 4:**
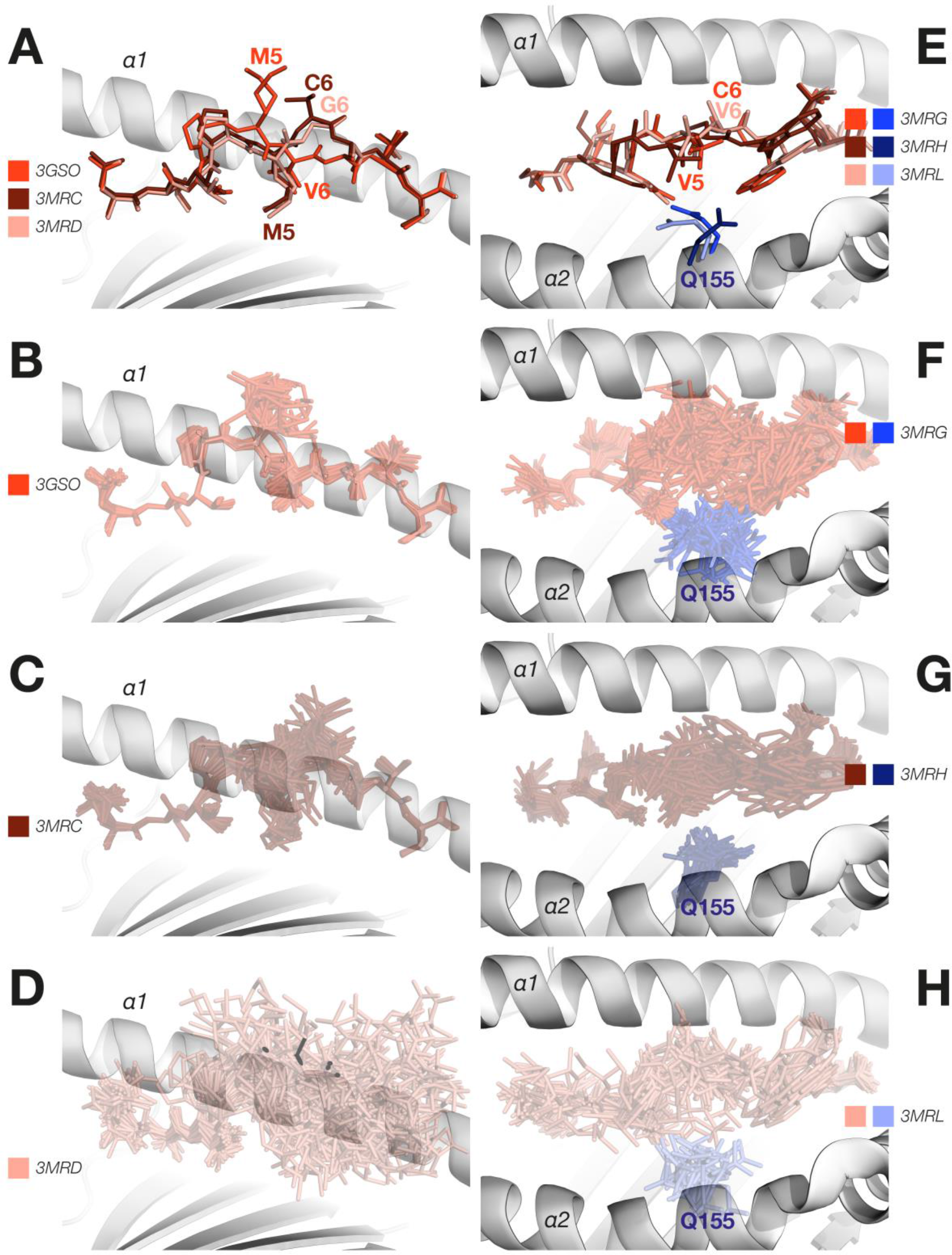
Prominence of MHC-bound peptides in TCR recognition. **A.** Original peptide conformations of HLA-A*02:01-NLV (red) (PDB ID 3GSO), HLA-A*02:01-NLV_V6C_ (brown) (PDB ID 3MRC), and HLA-A*02:01-NLV_V6G_ (pale red) (PDB ID 3MRD) in the original structures, compared to ensemble results for **B.** HLA-A*02:01-NLV, **C.** HLA-A*02:01-NLV_V6C_, and **D.** HLA-A*02:01-NLV_V6G_, showing that the variability within the NLV_V6G_ variant ensemble substantially exceeds the variation between any of the systems. The MHC α2 helix is omitted in all left-hand side panels. **E.** An overlay of the original peptide conformations of HLA-A*02:01-CIN (red/blue) (PDB ID 3MRG), HLA-A*02:01-CIN_N3S_ (brown/navy) (PDB ID 3MRH), and HLA-A*02:01-CIN_C6V_ (pale red/pale blue) (PDB ID 3MRL), compared to ensemble results for **F.** HLA-A*02:01-CIN, **G.** HLA-A*02:01-CINN3S and **H.** HLA-A*02:01-CIN_C6V_, showing that the variability within each ensemble is similar in magnitude to the variability between systems. The highly flexible P5 and P6 residues are found around the center of the peptide.

### Implications of conformational flexibility for vaccine design

Ensemble refinement also has practical applications in the analysis of pMHC systems for vaccine design. Previous research^18^ has sought to determine the structural basis for why the 2009 pandemic influenza virus, A/Auckland/1/2009(H1N1), exhibits a relatively high rate of cross-reactive CD8+ T cells with the 1918 strain A/Brevig Mission/1/1918(H1N1), but much lower cross-reactivity with seasonal strains from intermediate years (e.g. 1947 and 1972). Structures of HLA-B*35:01 complexed with six nucleoprotein (NP)418 peptides derived from influenza virus strains from six different years spanning the period 1918 to 2009 were determined and analyzed. The interpretation of these structures was that the solvent exposed P5-Arg residue common to both the 1918-NP418 (Figure 5A) and 2009-NP418 (Figure 5C) peptides produced a different surface recognition profile compared to the Asp-Lys (DK) motif found in NP418 peptides from intermediate years (e.g. 1972, Figure 5E), and that this may partly explain the differential CD8+ T cell cross-reactivity. In particular, it was suggested that the P5-Lys found in the NP418 of intermediate years (1947 and 1972) is too distal from the neighboring MHC residue Gln155 to allow the formation of the same hydrogen bond that is found between the P5-Arg and Gln155 residues in 1918-NP_418_ and 2009-NP_418_.

**Figure 5:**
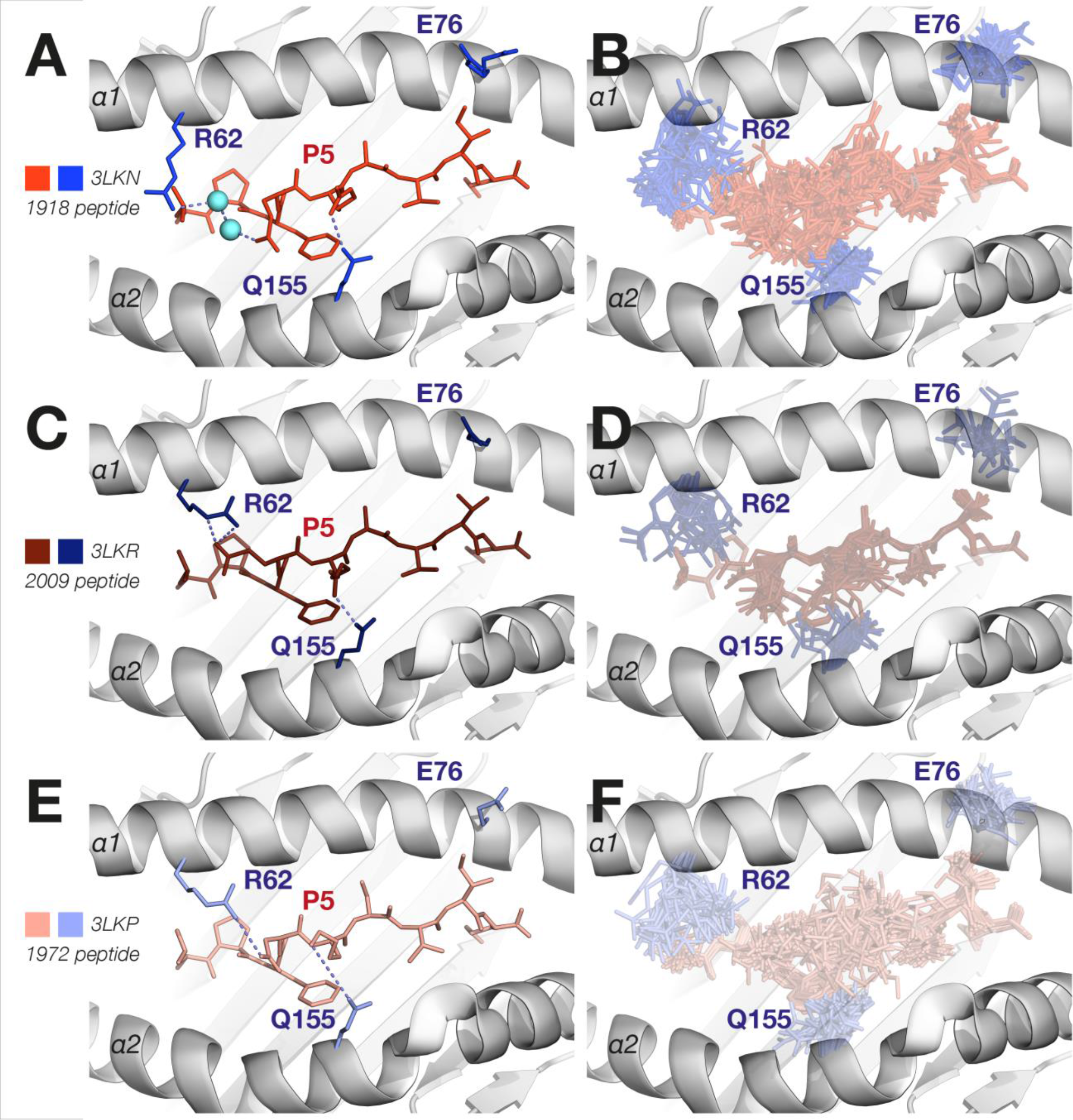
Implications of conformational flexibility for vaccine design. **A.** Original structure of 1918 influenza NP peptide bound to HLA-B*35:01 (PDB ID 3LKN) compared with **B.** ensemble results from the same system. The MHC residues Arg62, Glu76, and Gln155 are shown in blue sticks. **C.** Original structure of 1972 influenza NP peptide bound to HLA-B*35:01 (PDB ID 3LKP) compared with **D.** ensemble results from the same system. The MHC residues Arg62, Glu76, and Gln155 are shown in navy sticks. **E.** Original structure of 2009 influenza peptide bound to HLA-B*35:01 (PDB ID 3LKR) compared with **F.** ensemble results from the same system. The degree of flexibility of the 1918 and 1972 systems is seen to be similar in magnitude to the difference between the systems. The MHC residues Arg62, Glu76, and Gln155 are shown in pale blue sticks. Hydrogen bonds are shown as dashed lines.

Our ensemble results provide a new perspective on these structural differences by showing that the P5 residues for all three systems exhibit considerable flexibility, with hydrogen bonds formed with the Gln155 residue in some of the ensemble structures but not others (Figure 5B for 1918 strain, 5F for 1972 strain, and 5D for 2009 strain). We further explored the dynamics of HLA-B*35:01-1918-NP_418_ through MD, which showed even greater flexibility of the MHC Arg62 and Gln155 side chains than observed in the ensemble (Figure S1G and S1H). Both the ensemble and the MD results concur that the range of conformations adopted by the 1918-NP418 peptide when bound to HLA-B*35:01 is greater than any differences found between the static conformations of different years. The two distinct conformations of the Arg62 residue found in different years (compare Figure 5A and 5B), were also observed in all three of our ensembles. These results indicate that rather than focusing on identifying single differences between static structures, a richer and more complete description of influenza pMHC systems will analyze differences and similarities in the distribution of conformations found in different years. While the original study concluded that identifying key solvent-exposed residues might be important for vaccine design, our results highlight the critical importance of considering conformational dynamics of the peptide variants in the development of new vaccines, since conformational flexibility is a fundamental characteristic feature of these pMHC systems.

### TCR recognition models

By providing a means of examining the conformational diversity of pMHC systems, ensemble refinement has the potential to provide much richer information for discerning between induced fit and conformational selection models of TCR binding than is possible with traditional structure-based methods. For example, the structure of HLA-A*02:01 bound to the human T-lymphotropic virus type 1 (HTLV-1) Tax peptide was compared to the structure of HLA-A*02:01 bound to the similar Tel1p peptide from *Saccharomyces cerevisiae*^19^ (Figure 6A). Despite very similar conformations when bound to the MHC alone, the two peptides adopted substantially different conformations upon A6 TCR recognition, distinguishable by a ∼180° flip of the peptide backbone *ϕ* angle at P6, and different orientations of P4 and P5 side chains (Figure 6A). These structural observations were complemented with experimental measurements and MD, which suggested that these conformational differences are the result of the greater inherent flexibility of the Tel1p peptide relative to the Tax peptide, which enables the former to adopt a conformational binding pattern inaccessible to the latter. Our ensemble results provide strong support for this hypothesis, as both conformations of the Tel1p peptide, in particular the bond flip of P5, are present in the ensembles (Figure 6B). The structural variation found within the Tel1p ensemble is more than sufficient to encompass the alternate peptide conformation associated with the A6 TCR-bound form, thereby supporting the contention that the Tel1p conformation observed following A6 TCR binding is due to a conformational selection mechanism, rather than to induced fit^19^. This finding is further corroborated by our MD results, which show very similar bond-flipping behavior to that observed in the ensembles (Figure S1I and S1J).

**Figure 6:**
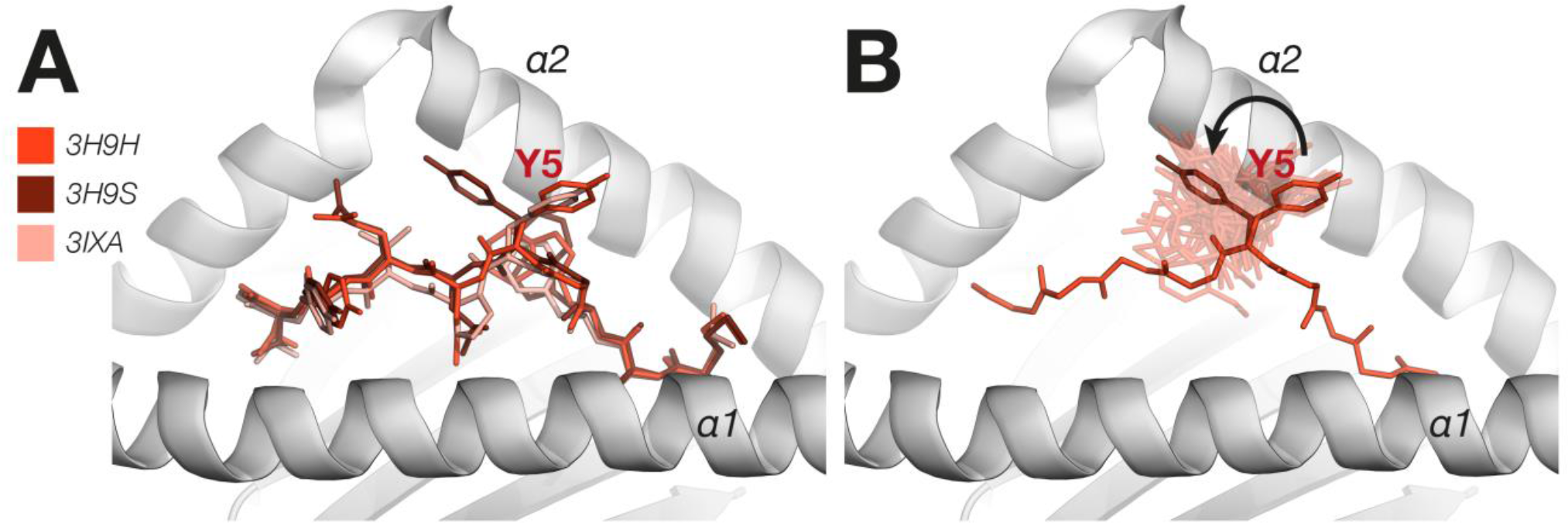
TCR recognition models. A. An overlay of HLA-A*02:01-Tel1p without A6 TCR (red) (PDB ID 3H9H), HLA-A*02:01-Tel1p with A6 TCR (brown) (PDB ID 3H9S), and HLA-A*02:01-Tax without the TCR (pale red) (PDB ID 3IXA) compared to B. the original HLA-A*02:01-Tel1p without A6 TCR (red) with overlaid ensemble results (transparent red). The bond flip of P5 distinguishing bound and unbound peptide forms is observed within the ensemble of the original Tel1p structure without TCR.

Another study concerning TCR recognition models examined the basis of LC13 TCR alloreactivity by comparing the structures and binding modes of two peptides (which are referred to as ‘allotope’ and ‘mimotope’) bound to three different HLA-B*44 alleles (HLA-B*44:02, HLA-B*44:03 and HLA-B*44:05)^20^. The purpose of the study was to understand the mechanism underpinning LC13 alloreactivity with peptide-bound HLA-B*44:02 and HLA-B*4405, but not HLA-B*44:03, which differs from the other alleles by 1 and 2 polymorphisms, respectively. The interpretation based on the static structures is that the HLA-B*44:05-bound mimotope and allotope peptides adopt a similar conformation only *after* LC13 TCR ligation, indicating an induced-fit ‘forcing’ by the TCR. Specifically, for the allotope a downward shift of the P3 residue upon binding to the TCR is observed (Figure 7A), while the mimotope exhibited an even more dramatic rearrangement of the P3, P5, and P7 side chains following TCR binding (Figure 7C). It is proposed that the reason the HLA-B*44:05 and HLA-B*44:02 complexes bind with the LC13 TCR, while the HLA-B*44:03 complex does not, is because the latter does not accommodate the conformational change to the peptide necessary for TCR recognition. Specifically, it is suggested that P3 of the allotope complex is being prevented from rotating into the TCR-binding position by the unfavorable location of Leu156 in HLA-B*44:03 (Figure 7E).

**Figure 7:**
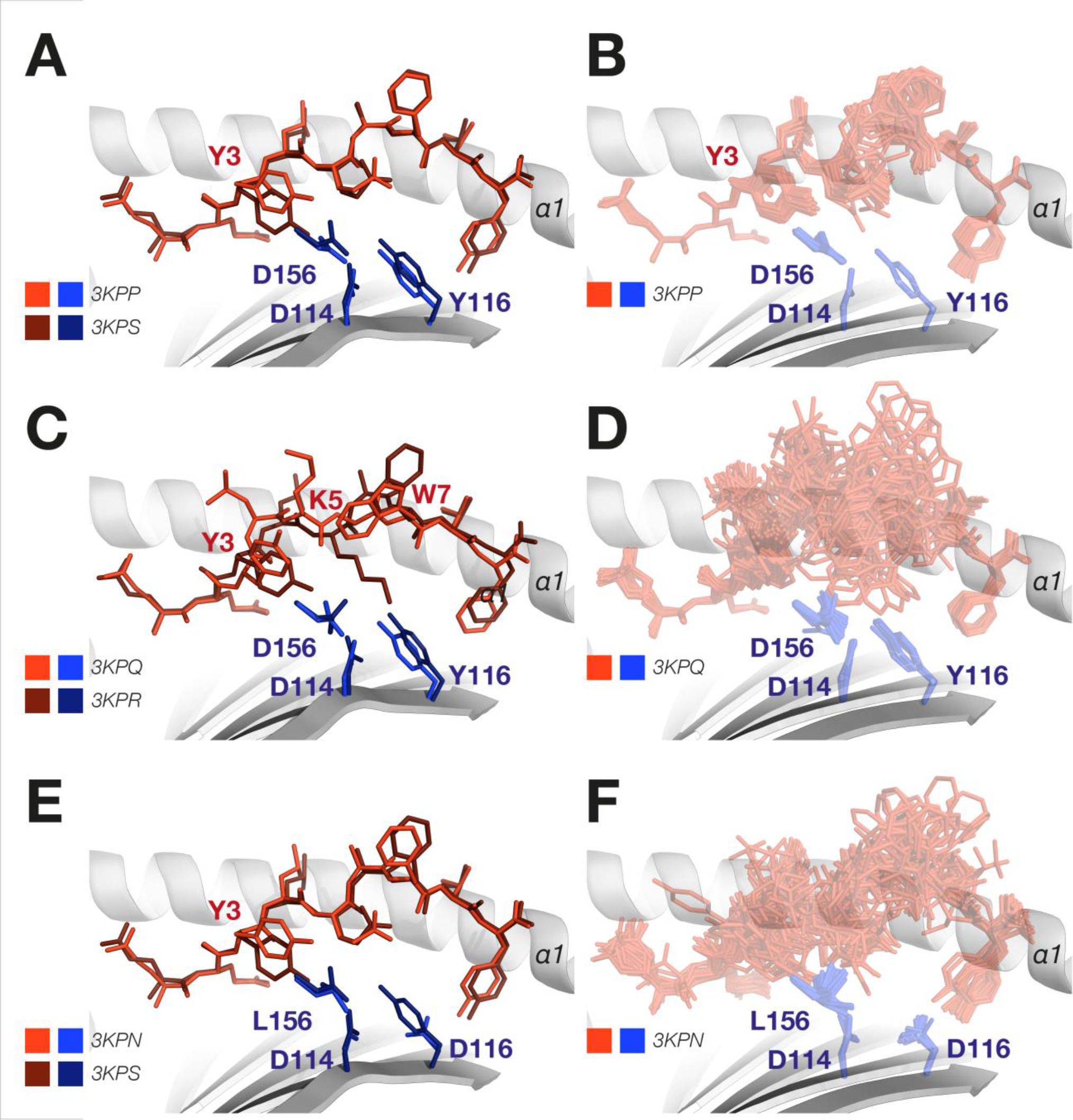
Conformational Plasticity and Molecular Mimicry. **A.** Overlay of the original structures of the HLA-B*44:05-allotope unbound (red/blue) (PDB ID 3KPP) and bound to TCR (brown/navy) (PDB ID 3KPS), compared to **B.** the ensemble results for HLA-B*44:05-allotope, showing that structural variability within the ensemble is greater than the difference between TCR-bound and unbound structures. The MHC α2-helices have been omitted. **C.** Overlay of the original structures of the HLA-B*44:05-mimotope unbound (red/blue) (PDB ID 3KPQ) and bound to TCR (brown/navy) (PDB ID 3KPR), compared to **D.** the ensemble results for the HLA-B*44:05-mimotope ensemble, showing that structural variability within the ensemble is greater than the difference between TCR-bound and unbound structures. The MHC α2-helices have been omitted. **E.** Overlay of the original structures of TCR-ligated HLA-B*44:05-allotope (brown/navy) (PDB ID 3KPS) with the unbound HLA-B*44:03-allotope (red/blue) (PDB ID 3KPN), showing the different position of P5 which is alleged to be the result of the unfavourable position of Leu156. **F.** The ensemble results for the HLA-B*44:03-allotope complex, however, show that the P5 residue is highly flexible, adopting a range of conformations overlapping with that found in the TCR-bound form of the HLA-B*44:05 complex. The MHC α2-helices have been omitted.

Our ensemble results provide new information pertinent to this analysis, since all HLA-B*44:05 allotope and mimotope complexes exhibit considerable flexibility (Figures 7B and 7D), with both the unbound and TCR-bound peptide conformations clearly present within each ensemble. These results indicate that, rather than being induced by the TCR, the conformational plasticity originally described may instead be due to pre-existing conformational variability with each system. The ensemble of the HLA-B*44:03 complex likewise exhibits considerable flexibility (Figure 7F), with the P5 residue clearly adopting a range of conformations, including the conformation seen in the TCR-bound forms of the HLA-B*44:05 system believed to be inaccessible for this allele. This result highlights how extracting the dynamical information already contained within the crystallographic data can further inform functional interpretations.

Overall our data highlights the fact that observed differences between static TCR-bound and unbound pMHC are not necessarily due to TCR binding, but rather the natural flexibility of each system. The traditional focus on static structures may have inadvertently had the effect of directing analyses in favour of an induced fit mechanism, since in analyzing a single structure the underlying conformational heterogeneity of the initial system which forms the basis for a conformational selection mechanism is overlooked.

## Discussion

X-ray crystallography has played a considerable role in increasing our understanding of the structural basis of TCR-pMHC engagement, but is limited in that it provides only a static snapshot. The role of conformational dynamics in TCR recognition of the pMHC has therefore tended to be overlooked, despite its important role in determining biological outcomes. Our study strongly emphasizes that a sizeable number of pMHC complexes exhibit considerable conformational flexibility that is not apparent in the published single-structure crystallographic model. Such dynamic variability, shown in both our ensemble and MD results, extends well beyond mere alternate side-chain rotamers or bond flips, and encompasses a vast range of motions including large-scale backbone movements and interconversion between substantively different peptide conformations. This result initially seems at odds with the relatively low B-factors reported in some of these systems. However, it is likely that B-factors substantially underestimate atomic motions, such that much of information on the flexibility and motions present in the molecule is lost in the refinement process^21, 22^. Our results corroborate these findings, and highlight the importance of using ensemble refinement techniques to more accurately gauge protein conformational flexibility.

Ensemble methods are subject to many of the same limitations as traditional single-structure methods. In particular, they are liable to be biased by crystal packing, artefacts of performing experiments at cryogenic temperatures, and over-fitting or other biases introduced in the refinement process. The distinctive advantage of ensemble methods is that they improve the usefulness and accuracy of structural models by extending them from showing features in only three dimensions, to also showing at least part of the ‘fourth dimension’ of how the structure varies over time. Biological macromolecules such as proteins exist out of thermodynamic equilibrium with their surroundings, and as such motion and variability over time is an intrinsic aspect of their behaviour. Ensemble methods recognize this by incorporating such variation as a key component of structural analysis, not merely as a minor complication or an afterthought during analysis.

Our study complements and extends the results of existing studies which have examined the dynamics of pMHC and TCR-pMHC systems using alternative methodologies. Previous MD studies of such systems have uncovered a similarly diverse range of conformations (especially of the peptide) for a number of different MHC alleles^4, 23^. Hydrogen-deuterium exchange has also been used to probe the dynamics at the TCR-pMHC interface, with a number of studies finding evidence of conformational flexibility which varies according to the MHC allele^24^, the bound peptide^25^, and upon TCR ligation^26^. The study of pMHC dynamics using nuclear magnetic resonance (NMR) spectroscopy is complicated by the relatively large size of these systems^27^, however several recent studies have all found evidence of significant conformational variation in both the peptide and MHC^28, 29^. The unique advantage of ensemble refinement relative to these existing techniques is that it enables direct visualization of conformational variation derived from already-existing crystallographic data, without the need to carry out additional experimental work or conduct lengthy MD simulations.

Many contemporary TCR-pMHC structural studies attempt to determine the factors that influence TCR binding specificity or cross-reactivity by comparing the structures of two or more complexes and identifying their similarities and differences. Thus, for instance, if a peptide adopts one conformation when bound to MHC alone and a different conformation when bound to MHC complexed with TCR, it is often inferred that this represents a change in conformation brought about by the binding of the TCR. However, as we have emphasized in the case studies discussed above, a baseline degree of structural heterogeneity must be expected in any protein system, resulting from the intrinsic flexibility of protein molecules. The precise position of backbone atoms, orientation of side chains, extent of hydrogen bonding networks, etc., is therefore expected to vary over time and between samples. Crystallographic structures are thus properly regarded as the result of random draws from the underlying distribution of protein conformational states. Observed structural differences between systems should therefore be compared in magnitude to the intrinsic structural variation within each system, to determine whether there is sufficient reason to justify claiming a significant difference between the systems. In principle, this is much the same as statistical tests routinely applied in disciplines outside of structural biology, where a difference between two groups is only deemed significant if this inter-group difference is sufficiently greater than the intra-group variation. Although it is possible to estimate coordinate error from the crystallographic data, no such method yet exists for statistical comparison of protein ensembles. The resulting failure to consider this baseline of structural heterogeneity has resulted in numerous false positive identifications of putative differences, and thereby hampered the growth in our understanding of MHC and T-cell function.

One possible method of statistical comparison would be to use a force field to compute the mean and standard deviation of energies in the ensemble, as well as the energy of a comparison conformation of interest, thereby enabling a Z-score to be calculated. While rigorous hypothesis testing is not possible since these energies cannot be assumed to be normally distributed, nevertheless low Z-scores (of less than one or two) would be an indication that the comparison conformation is well inside the range of energies found within the ensemble. In combination with visual inspection of the structural differences of the ensemble and comparison conformation, this crude test could be useful in quickly estimating whether the comparison conformation likely derives from a distinct energy landscape, or is simply part of the initial ensemble of states.

In addition to the question of whether two protein systems exhibit a structural difference that is significantly greater than background heterogeneity, there is the further question as to what is the most likely cause of this difference. In many structural studies it is inferred that any conformational differences between a TCR-bound and unbound pMHC complex have been induced by the TCR. Though this is undoubtedly the case for some systems, in other systems it is likely that causation runs in the opposite direction, meaning that the reason the TCR has bound to the peptide is because the peptide has adopted that particular conformation. This distinction corresponds to the dispute between the ‘induced fit’ and ‘conformational selection from existing equilibrium’ models of protein interaction. As we noted previously, ensemble refinement provides a method uniquely suited to examine this question, since if the TCR-bound peptide conformation is already observed in the unbound pMHC ensemble, this constitutes at least *prima facie* evidence for a conformational selection mechanism^30^. Our ensemble results show that, at least for some peptide-MHC systems, there is a considerable degree of intrinsic flexibility from which the TCR has considerable ‘choice’ in selecting a favorable form with which to bind. Recent work in support of these ideas has shown that conformational variability may facilitate several rounds of association and disassociation of peptides with MHC complexes, with low affinity peptides rapidly disassociating and thereby gradually establishing an equilibrium ensemble where a given MHC is only bound to its corresponding high affinity peptides^31^. Our results build upon these findings in providing further evidence of the importance of a structural ensemble-based approach in the study of MHC dynamics and affinity.

A third, related and important consideration when examining TCR-pMHC systems is that the intrinsic structural heterogeneity resulting from peptide and MHC flexibility need not always be simply random fluctuations about a single functional state. Rather, it is likely that in at least some cases different conformations of the system fulfil different functional purposes. As discussed in a number of the cases we considered, the flexibility of the system itself can play a role in determining binding specificity, for example through bond flips that alternate between bound and unbound peptide conformations. Such behaviors indicate that one cannot simply infer that because a peptide is observed in one particular conformation when crystallized bound to the TCR, this was the same conformation that the peptide adopted during the process of TCR recognition — it is possible that conformational changes may occur *after* the initial binding event. There may be one or more intermediates and/or transition states with subtly or considerably different conformations to that seen in the final complex, with the conformational variability an intrinsic part of the binding process^29^. Peptide flexibility should therefore be considered from the outset as an essential functional aspect of the system when analyzing peptide, MHC and TCR binding dynamics.

## Conclusions

Our analysis highlights the breadth of insights related to TCR recognition of pMHC that can be gained by using ensemble refinement to extract the wealth of dynamical information available in existing crystallographic data, information that is overlooked in the traditional practice of analyzing single isolated structures. Examination of a variety of TCR-pMHC complexes suggests that reliance on single structures entails significant limitations for understanding the rules of productive TCR ligation, particularly for static interpretations involving fine details such as hydrogen-bonding networks and side chain orientation. Furthermore, our findings indicate that structural differences between the pMHC and TCR-pMHC conformations may be due to intrinsic flexibility rather than any change elicited by binding itself, and highlight the risk of false-positives in assuming that an observed difference between static pMHC and TCR-pMHC complexes is necessarily due to the binding process, rather than a reflection of the natural flexibility of each system. Broadly, our results indicate that explaining biological function with reference to the properties of protein ensembles will lead to a more accurate and comprehensive understanding of underlying mechanisms. As such, we propose that extracting dynamic information from X-ray data through the use of ensemble methods will lead to a richer, dynamic understanding of TCR-pMHC interactions.

## Materials and Methods

### Computational resources

Calculations, modeling and simulations were performed on a range of computing resources: ORCHARD 800 core x86 cluster (Monash University; X-ray ensemble refinement); Multi-modal Australian ScienceS Imaging and Visualisation Environment (MASSIVE; atomistic MD).

### Ensemble refinement

All atomic coordinates (.pdb) and crystallographic reflections (.mtz) files were sourced from the PDB_REDO server^32^. Ensembles were calculated with PHENIX 1.9-1692^33^ by first passing each system through the ReadySet tool, and then using the Ensemble refinement tool with default parameters^8^. Ensemble analysis was performed using PyMOL version 1.3^34^ and python scripts.

### Atomic coordinates, modeling and graphics

In MD simulations, atomic coordinates were obtained from the following PDB entries: 1UXS, 2A83, 3H9H, 2LKN, and 3MRD. Structural representations were produced using PyMOL version 1.3^34^ and VMD 1.9.1^35^. Trajectory manipulation and analysis was performed using MDTraj^36^ and VMD 1.9.1^35^.

### Molecular dynamics (MD) systems setup and simulation

Each protein, with protonation states appropriate for pH 7.0^37, 38^, was placed in a rectangular box with a border of at least 12 Å, explicitly solvated with TIP3P water^39^, counter-ions added, and parameterized using the AMBER ff14SB all-atom force field^14,40,41^. After an energy minimization stage and an equilibration stage, production simulations were performed in the NPT ensemble. Four independent replicates of each system were simulated for 500 ns each using NAMD 2.9^42^.

## Acknowledgements

This work was supported by grants from the National Health and Medical Research Council (NHMRC) and the Australian Research Council (ARC). NAB is funded by an ARC Future Fellowship (110100223). This work was supported by the Multi-modal Australian ScienceS Imaging and Visualisation Environment (MASSIVE) (www.massive.org.au).

## Author Contributions

AMB designed the study. JF performed ensemble refinement and molecular dynamics simulations. BTR prepared the figures. JF, BTR, NAB and AMB wrote the manuscript.

## Competing financial interests

The authors declare no competing financial interests.

## Supporting Information

**Figure S1:**
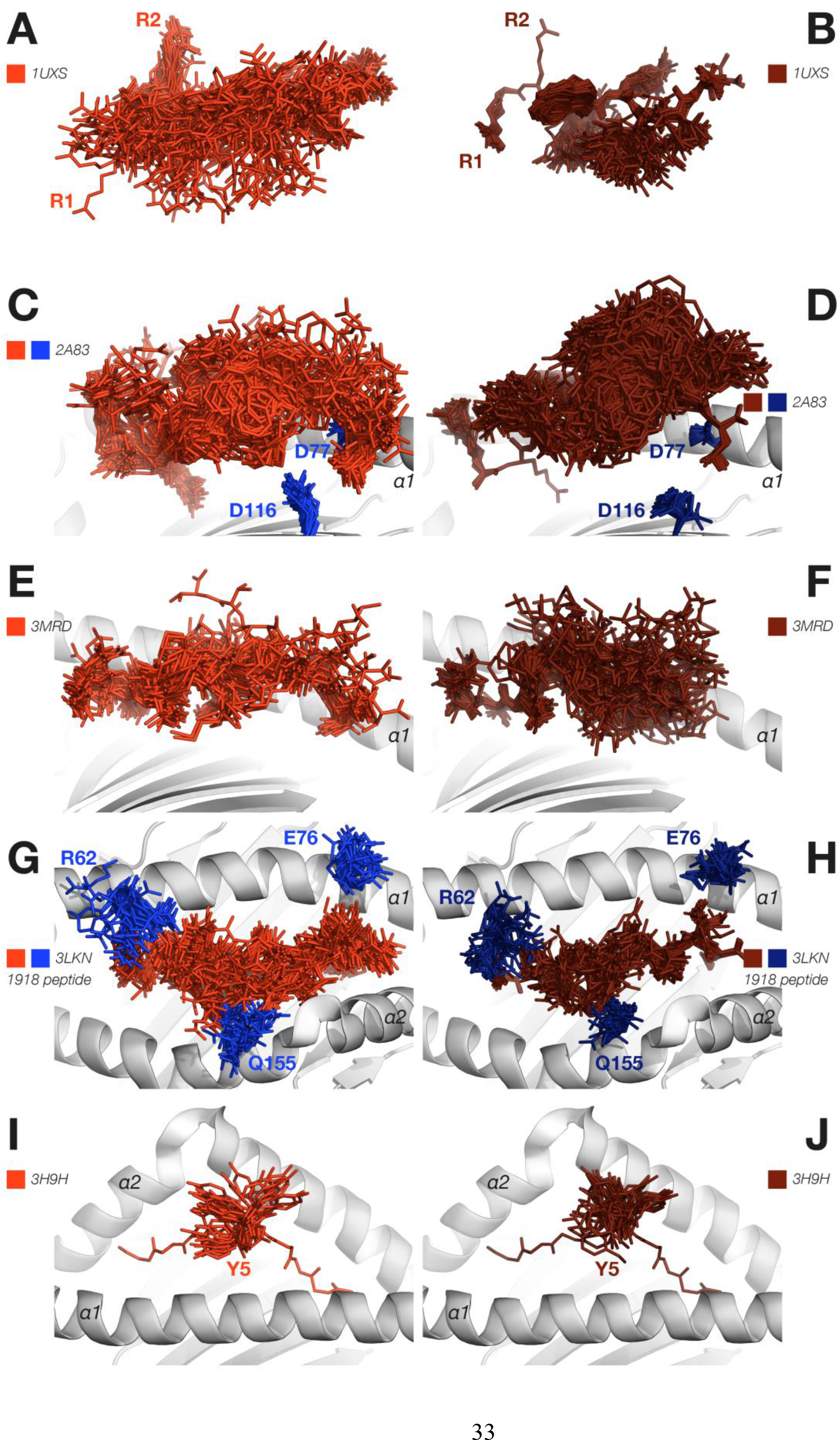
Molecular Dynamics and Ensemble Refinement Comparison. **A.** Overlay of 50 frames from 2 μs MD simulation of an Epstein-Barr virus peptide pLMP2 bound to HLA-B*27:09 (red sticks) (PDB ID 1UXS) compared to **B.** ensemble results of the same system (brown sticks), showing very similar patterns of dynamics. MHC α-helices have been omitted. **C.** Overlay of 50 frames from 2 μs MD simulation of a glucagon-derived self-peptide (pGR) (red sticks) bound to HLA-B*27:09 (blue sticks, cartoon) (PDB ID 2A83) compared to **D.** ensemble results of the same system (peptide as brown sticks; MHC as navy sticks, cartoon), with Asp77 and Asp116 shown as sticks, showing very similar patterns of dynamics. The MHC α2-helix has been omitted. **E.** Overlay of 50 frames from 2 μs MD simulation of NLV_V6G_ peptide (red sticks) bound to HLA-A*02:01 (cartoon) (PDB ID 3MRD) compared to **F.** ensemble results of the same system (peptide as brown sticks; MHC as cartoon), showing comparable overall degrees of flexibility. The MHC α2 helix has been omitted. **G.** Overlay of 50 frames from 2 μs MD simulation of 1918-NP_418_ (red sticks) bound to HLA-B*35:01 (blue sticks, cartoon) (PDB ID 3LKN) compared to **H.** ensemble results of the same system (peptide as brown sticks; MHC as navy sticks, cartoon), with very similar patterns of flexibility evident in each. The MHC residues Arg62, Glu76, and Gln155 are shown as sticks. **I.** Overlay of 50 frames from 2 μs MD simulation of Tel1p (red sticks) bound to HLA-A*02:01 (cartoon) (PDB ID 3H9H) compared to **J.** ensemble results of the same system (peptide as brown sticks; MHC as cartoon), showing the very similar bond flips of P5-Tyr in both cases.

## References

1 van den Bedem, H. & Fraser, J. S. Integrative, dynamic structural biology at atomic resolution [mdash] it’s about time. Nature methods 12, 307–318 (2015).

2 Woldeyes, R. A., Sivak, D. A. & Fraser, J. S. E pluribus unum, no more: from one crystal, many conformations. Current opinion in structural biology 28, 56–62 (2014).

3 Furnham, N., Blundell, T. L., DePristo, M. A. & Terwilliger, T. C. Is one solution good enough? Nature structural & molecular biology 13, 184–185 (2006).

4 Kass, I., Buckle, A. M. & Borg, N. A. Understanding the structural dynamics of TCR-pMHC interactions. Trends in immunology 35, 604–612 (2014).

5 Lange, O. F., Van der Spoel, D. & De Groot, B. L. Scrutinizing molecular mechanics force fields on the submicrosecond timescale with NMR data. Biophysical journal 99, 647–655 (2010).

6 Lindorff-Larsen, K. et al. Systematic validation of protein force fields against experimental data. PloS one 7, e32131 (2012).

7 Piana, S., Lindorff-Larsen, K. & Shaw, D. E. How robust are protein folding simulations with respect to force field parameterization? Biophysical journal 100, L47–L49 (2011).

8 Burnley, B. T., Afonine, P. V., Adams, P. D. & Gros, P. Modelling dynamics in protein crystal structures by ensemble refinement. Elife 1, e00311 (2012).

9 Correy, G. J. et al. Mapping the accessible conformational landscape of an insect carboxylesterase using conformational ensemble analysis and kinetic crystallography. Structure 24, 977–987 (2016).

10 Forneris, F., Burnley, B. T. & Gros, P. Ensemble refinement shows conformational flexibility in crystal structures of human complement factor D. Acta Crystallographica Section D: Biological Crystallography 70, 733–743 (2014).

11 Chen, J.-L. et al. Ca(2+) release from the endoplasmic reticulum of NY-ESO-1 specific T cells is modulated by the affinity of T cell receptor and by the use of the CD8 co-receptor. Journal of immunology (Baltimore, Md. : 1950) 184, 1829–1839, doi:10.4049/jimmunol.0902103 (2010).

12 Borbulevych, O. Y., Baxter, T. K., Yu, Z., Restifo, N. P. & Baker, B. M. Increased immunogenicity of an anchor-modified tumor-associated antigen is due to the enhanced stability of the peptide/MHC complex: implications for vaccine design. The Journal of Immunology 174, 4812–4820 (2005).

13 Fiorillo, M. T. et al. Allele-dependent Similarity between Viral and Self-peptide Presentation by HLA-B27 Subtypes. Journal of Biological Chemistry 280, 2962–2971, doi:10.1074/jbc.M410807200 (2005).

14 Maier, J. A. et al. ff14SB: Improving the Accuracy of Protein Side Chain and Backbone Parameters from ff99SB. Journal of Chemical Theory and Computation 11, 3696–3713 (2015).

15 Rückert, C. et al. Conformational dimorphism of self-peptides and molecular mimicry in a disease-associated HLA-B27 subtype. Journal of Biological Chemistry 281, 2306–2316 (2006).

16 Miles, J. J. et al. TCRα Genes Direct MHC Restriction in the Potent Human T Cell Response to a Class I-Bound Viral Epitope. The Journal of Immunology 177, 6804–6814, doi:10.4049/jimmunol.177.10.6804 (2006).

17 Reiser, J.-B. et al. Analysis of Relationships between Peptide/MHC Structural Features and Naive T Cell Frequency in Humans. The Journal of Immunology 193, 5816–5826, doi:10.4049/jimmunol.1303084 (2014).

18 Gras, S. et al. Cross-reactive CD8(+) T-cell immunity between the pandemic H1N1-2009 and H1N1-1918 influenza A viruses. Proceedings of the National Academy of Sciences of the United States of America 107, 12599–12604, doi:10.1073/pnas.1007270107 (2010).

19 Borbulevych, O. Y. et al. T cell receptor cross-reactivity directed by antigen-dependent tuning of peptide-MHC molecular flexibility. Immunity 31, 885–896 (2009).

20 Macdonald, W. A. et al. T Cell Allorecognition via Molecular Mimicry. Immunity 31, 897–908, doi:10.1016/j.immuni.2009.09.025.

21 Kuzmanic, A., Pannu, N. S. & Zagrovic, B. X-ray refinement significantly underestimates the level of microscopic heterogeneity in biomolecular crystals. Nature communications 5 (2014).

22 García, A. E., Krumhansl, J. A. & Frauenfelder, H. Variations on a theme by Debye and Waller: from simple crystals to proteins. Proteins: Structure, Function, and Bioinformatics 29, 153–160 (1997).

23 Narzi, D. et al. Dynamical characterization of two differentially disease associated MHC class I proteins in complex with viral and self-peptides. Journal of molecular biology 415, 429–442 (2012).

24 Fabian, H. et al. HLA-B27 subtypes differentially associated with disease exhibit conformational differences in solution. Journal of molecular biology 376, 798–810 (2008).

25 Hawse, W. F. et al. Peptide modulation of class I major histocompatibility complex protein molecular flexibility and the implications for immune recognition. Journal of Biological Chemistry 288, 24372–24381 (2013).

26 Hawse, W. F. et al. Cutting edge: evidence for a dynamically driven T cell signaling mechanism. The Journal of Immunology 188, 5819–5823 (2012).

27 Insaidoo, F. K., Zajicek, J. & Baker, B. M. A general and efficient approach for NMR studies of peptide dynamics in class I MHC peptide binding grooves. Biochemistry 48, 9708–9710 (2009).

28 Hawse, W. F. et al. TCR scanning of peptide/MHC through complementary matching of receptor and ligand molecular flexibility. The Journal of Immunology 192, 2885–2891 (2014).

29 Yanaka, S. & Sugase, K. Exploration of the Conformational Dynamics of Major Histocompatibility Complex Molecules. Frontiers in Immunology 8 (2017).

30 Vogt, A. D. & Di Cera, E. Conformational selection is a dominant mechanism of ligand binding. Biochemistry 52, 5723–5729 (2013).

31 Garstka, M. A. et al. The first step of peptide selection in antigen presentation by MHC class I molecules. Proceedings of the National Academy of Sciences 112, 1505–1510, doi:10.1073/pnas.1416543112 (2015).

32 Joosten, R. P., Long, F., Murshudov, G. N. & Perrakis, A. The PDB_REDO server for macromolecular structure model optimization. IUCrJ 1, 213–220 (2014).

33 Adams, P. D. et al. PHENIX: a comprehensive Python-based system for macromolecular structure solution. Acta Crystallographica Section D: Biological Crystallography 66, 213–221 (2010).

34 Schrödinger, L. The PyMOL Molecular Graphics System, Version 1.3.

35 Humphrey, W., Dalke, A. & Schulten, K. VMD: visual molecular dynamics. Journal of molecular graphics 14, 33–38, 27-38 (1996).

36 McGibbon, R. T. et al. MDTraj: A Modern Open Library for the Analysis of Molecular Dynamics Trajectories. Biophysical journal 109, 1528–1532, doi:10.1016/j.bpj.2015.08.015 (2015).

37 Dolinsky, T. J., Nielsen, J. E., McCammon, J. A. & Baker, N. A. PDB2PQR: an automated pipeline for the setup of Poisson–Boltzmann electrostatics calculations. Nucleic acids research 32, W665–W667 (2004).

38 Søndergaard, C. R., Olsson, M. H., Rostkowski, M. & Jensen, J. H. Improved treatment of ligands and coupling effects in empirical calculation and rationalization of p K a values. Journal of Chemical Theory and Computation 7, 2284–2295 (2011).

39 Jorgensen, W. L., Chandrasekhar, J., Madura, J. D., Impey, R. W. & Klein, M. L. Comparison of simple potential functions for simulating liquid water. The Journal of Chemical Physics 79, 926–935, doi:http://dx.doi.org/10.1063/1.445869 (1983).

40 Joung, I. S. & Cheatham, T. E. Determination of Alkali and Halide Monovalent Ion Parameters for Use in Explicitly Solvated Biomolecular Simulations. The Journal of Physical Chemistry B 112, 9020–9041 (2008).

41 Li, P., Roberts, B. P., Chakravorty, D. K. & Merz Jr, K. M. Rational design of particle mesh Ewald compatible Lennard-Jones parameters for +2 metal cations in explicit solvent. Journal of chemical theory and computation 9, 2733–2748 (2013).

42 Phillips, J. C. et al. Scalable molecular dynamics with NAMD. Journal of computational chemistry 26, 1781–1802 (2005).

